## Supplementary figures and images for "Aberrant cohesin function in *Saccharomyces cerevisiae* activates Mcd1 degradation to promote cell lethality"

### S1 Fig

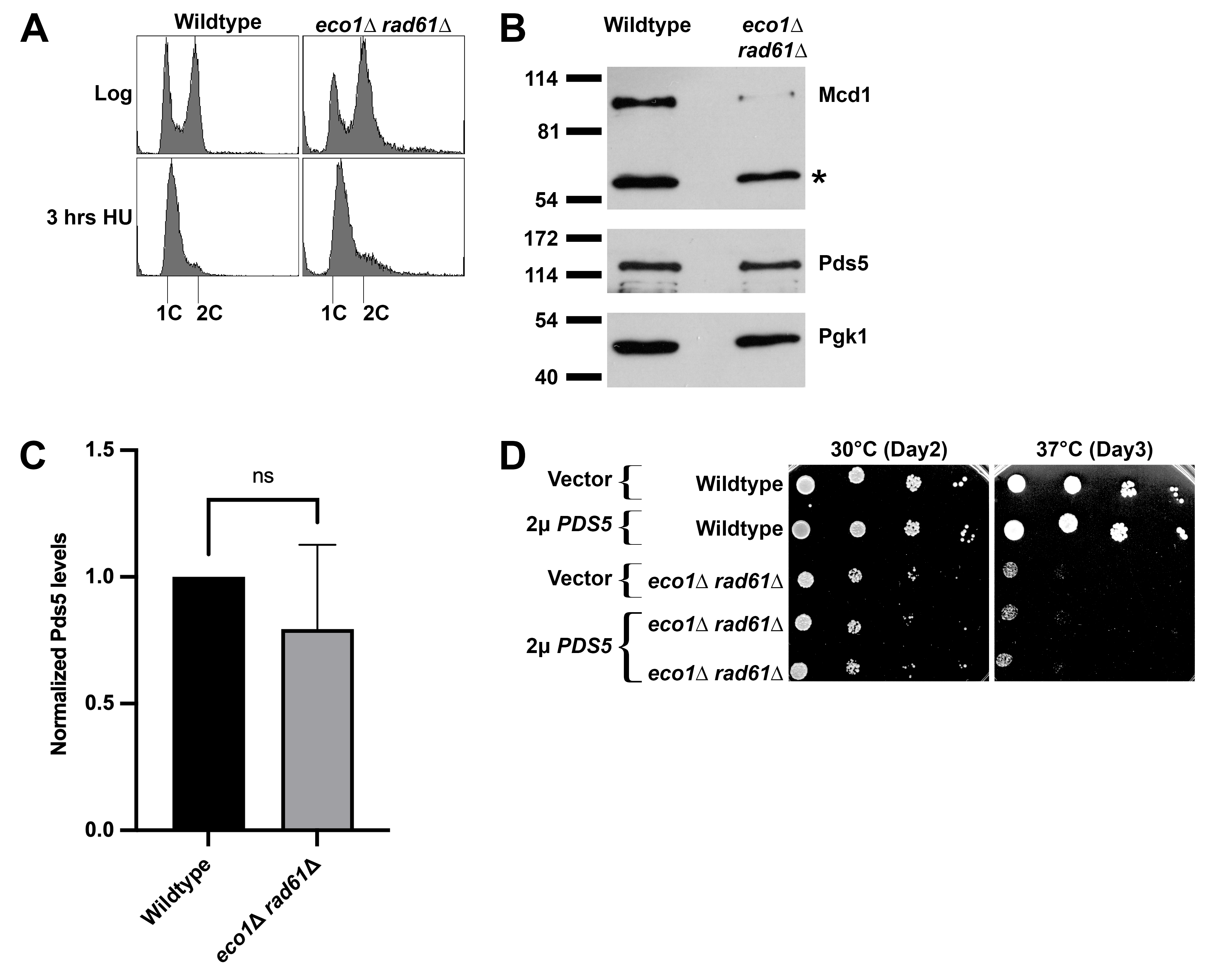

### S2 Fig

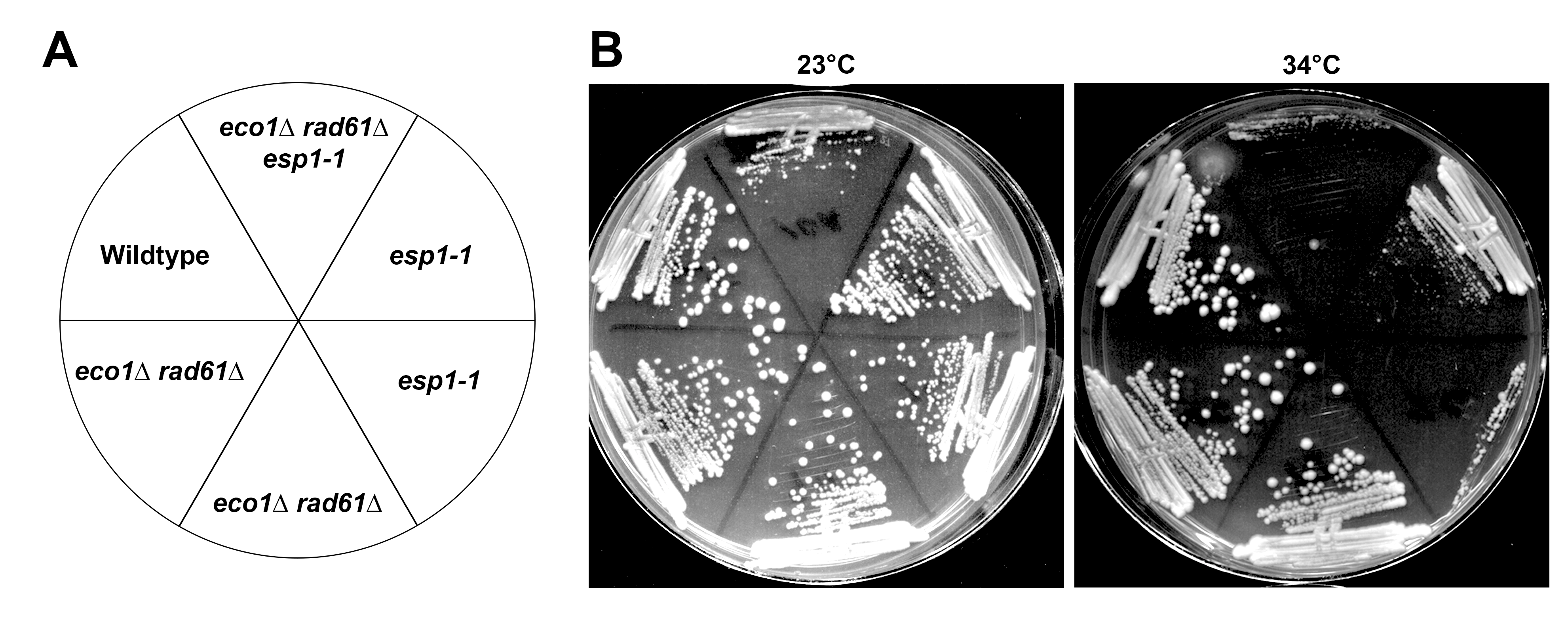

### S3 Fig

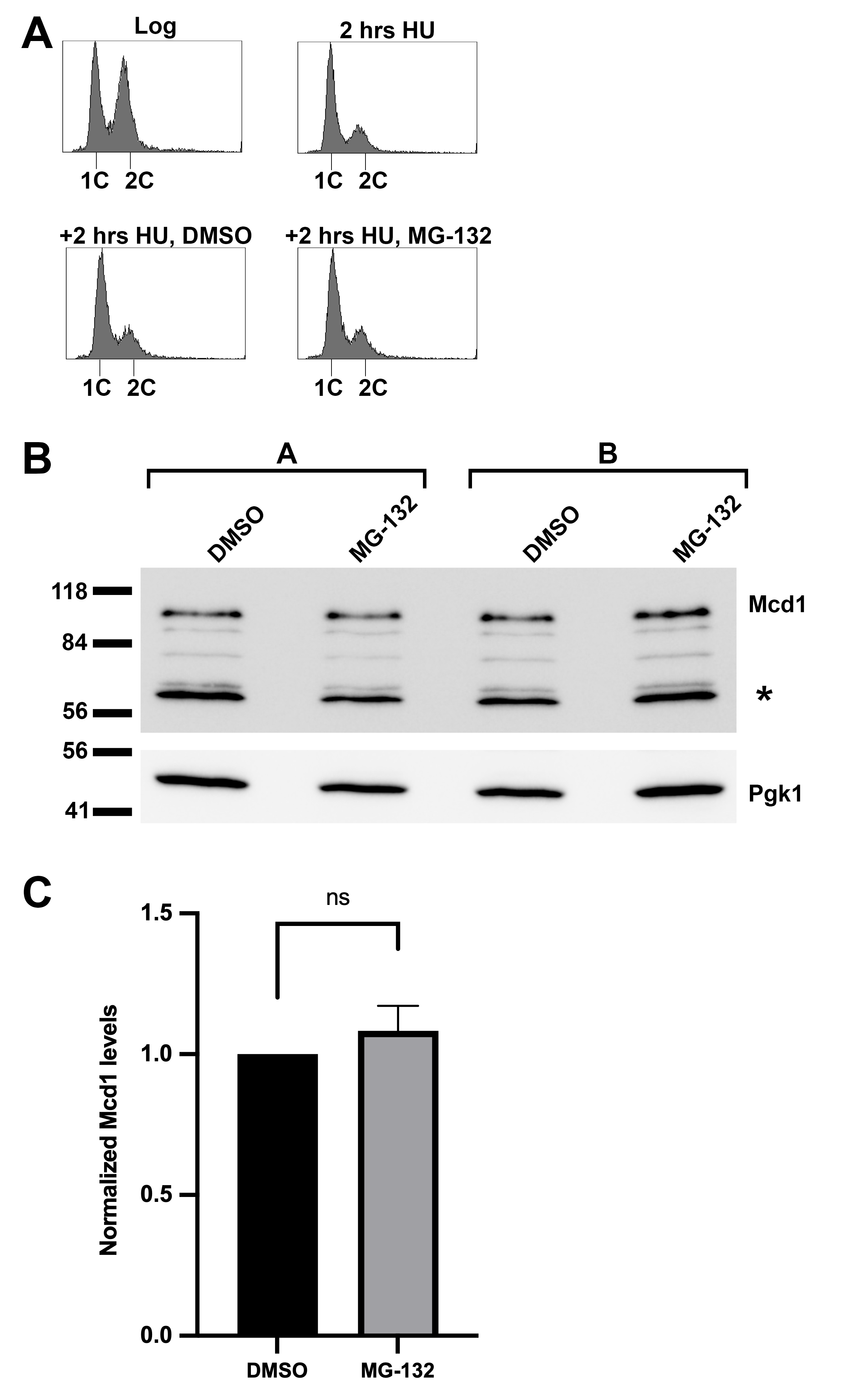

### S4 Fig

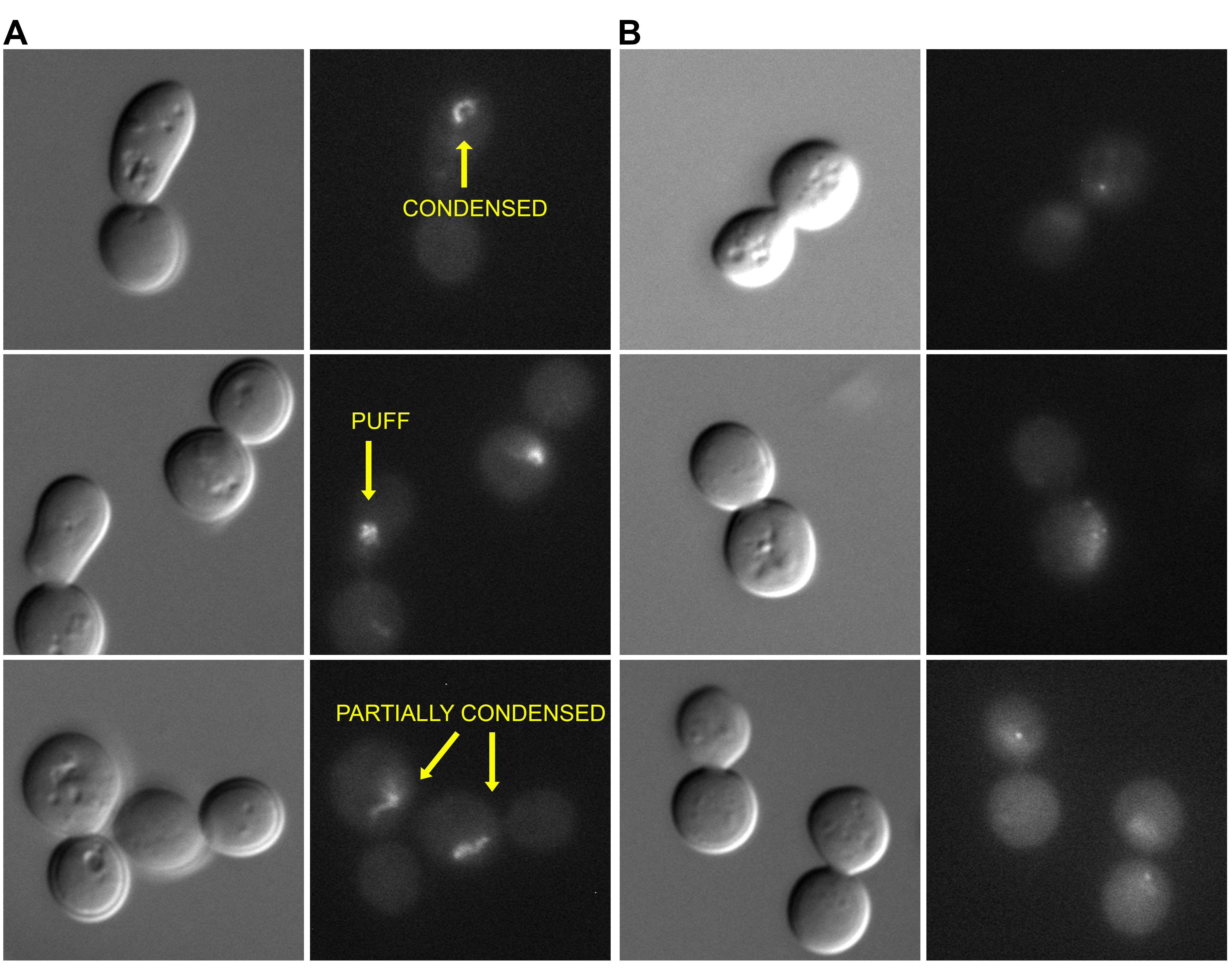
