## Supplementary material for "Aberrant cohesin function in *Saccharomyces cerevisiae* activates Mcd1 degradation to promote cell lethality": S1 Table

| Strain | Genotype | Reference |
|---|---|---|
| YPH499 | *MATa ura3-52 lys2-801_amber ade2-101_ochre trp1-Δ63 his3-Δ200 leu2-Δ1* | Sikorski and Hieter, 1989 |
| YBS514 | *MATa ura3-52 lys2-801 ade2-101 trp1-Δ63 his3-Δ200 leu2-Δ1 ctf7Δ1::HIS3 ctf7-203:LEU2* | Skibbens, 1999 |
| YMM808 | *MATa rad61::URA3 ade2-101 his3Δ200 leu2Δ1 lys2-801 trp1Δ63 ura3-52* | Maradeo and Skibbens, 2010 |
| YMM829 | *MATa eco1::HIS3 rad61::URA3 ade2-101 his3Δ200 leu2Δ1 lys2-801 trp1Δ63 ura3-52* | Maradeo and Skibbens, 2010 |
| YMM828 | *MATα eco1::HIS3 rad61::URA3 ade2-101 his3Δ200 leu2Δ1 lys2-801 trp1Δ63 ura3-52* | Maradeo and Skibbens, 2010 |
| YGS277 | *MATa bul2::TRP1 ura3-52 lys2-801_amber ade2-101_ochre trp1-Δ63 his3-Δ200 leu2-Δ1 (Isolate 1.1)* | This study |
| YGS279 | *MATa eco1::HIS3 rad61::URA3 bul2::TRP1 ade2-101 his3Δ200 leu2Δ1 lys2-801 trp1Δ63 ura3-52 (Isolate2A)* | This study |
| YGS280 | *MATa eco1::HIS3 rad61::URA3 bul2::TRP1 ade2-101 his3Δ200 leu2Δ1 lys2-801 trp1Δ63 ura3-52 (isolate2.1)* | This study |
| YGS309 | *MATa bre1::TRP1 ura3-52 lys2-801_amber ade2-101_ochre trp1-Δ63 his3-Δ200 leu2-Δ1 (isolate b.1)* | This study |
| YGS292 | *MATa eco1::HIS3 rad61::URA3 bre1::TRP1 ade2-101 his3Δ200 leu2Δ1 lys2-801 trp1Δ63 ura3-52 (Isolate 12.1)* | This study |
| YGS293 | *MATa eco1::HIS3 rad61::URA3 bre1::TRP1 ade2-101 his3Δ200 leu2Δ1 lys2-801 trp1Δ63 ura3-52 (Isolate 12.2)* | This study |
| YGS281 | *MATa ldb19::TRP1 ura3-52 lys2-801_amber ade2-101_ochre trp1-Δ63 his3-Δ200 leu2-Δ1 (Isolate 3.1)* | This study |
| YGS282 | *MATa eco1::HIS3 rad61::URA3 ldb19::TRP1 ade2-101 his3Δ200 leu2Δ1 lys2-801 trp1Δ63 ura3-52 (Isolate 4a)* | This study |
| YGS283 | *MATa eco1::HIS3 rad61::URA3 ldb19::TRP1 ade2-101 his3Δ200 leu2Δ1 lys2-801 trp1Δ63 ura3-52 (Isolate 4.2)* | This study |
| YGS284 | *MATa das1::TRP1 ura3-52 lys2-801_amber ade2-101_ochre trp1-Δ63 his3-Δ200 leu2-Δ1 (Isolate 5.1)* | This study |
| YGS286 | *MATa eco1::HIS3 rad61::URA3 das1::TRP1 ade2-101 his3Δ200 leu2Δ1 lys2-801 trp1Δ63 ura3-52 (Isolate 6.1)* | This study |
| YGS287 | *MATa eco1::HIS3 rad61::URA3 das1::TRP1 ade2-101 his3Δ200 leu2Δ1 lys2-801 trp1Δ63 ura3-52 (Isolate 6.3)* | This study |

| | | |
|---|---|---|
| YGS288 | *MATa san1::TRP1 ura3-52 lys2-801_amber ade2-101_ochre trp1-Δ63 his3-Δ200 leu2-Δ1 (Isolate 7.1)* | This study |
| YGS290 | *MATa eco1::HIS3 rad61::URA3 san1::TRP1 ade2-101 his3Δ200 leu2Δ1 lys2-801 trp1Δ63 ura3-52 (Isolate 8.2)* | This study |
| YGS291 | *MATa eco1::HIS3 rad61::URA3 san1::TRP1 ade2-101 his3Δ200 leu2Δ1 lys2-801 trp1Δ63 ura3-52 (Isolate 8.3)* | This study |
| YBS3164/ K699 | *MATa ade2-1 trp1-1 can1-100 leu2-3,112 his3-11,15 ura3 GAL psi+* | Gift from Dr. Guacci |
| YGS209 | *MATa ade2-1 trp1-1 can1-100 leu2-3,112 his3-11,15 ura3 GAL psi+ (2μm-TRP1)* | This study |
| YGS210 | *MATa ade2-1 trp1-1 can1-100 leu2-3,112 his3-11,15 ura3 GAL psi+ (2μm TRP1 MCD1)* | This study |
| YBS3170/ K5824 | *MATa smc3-42 ade2-1 trp1-1 can1-100 leu2-3,112 his3-11,15 ura3 GAL psi+* | Gift from Dr. Guacci |
| YGS229 | *MATa smc3-42 ade2-1 trp1-1 can1-100 leu2-3,112 his3-11,15 ura3 GAL psi+ (2μm-TRP1)* | This study |
| YGS230 | *MATa smc3-42 ade2-1 trp1-1 can1-100 leu2-3,112 his3-11,15 ura3 GAL psi+ (2μm TRP1 MCD1) (Isolate 6A)* | This study |
| YGS231 | *MATa smc3-42 ade2-1 trp1-1 can1-100 leu2-3,112 his3-11,15 ura3 GAL psi+ (2μm TRP MCD1) (Isolate 6B)* | This study |
| YBS3168/ K6013 | *MATa smc1-259 ade2-1 trp1-1 can1-100 leu2-3,112 his3-11,15 ura3 GAL psi+* | Gift from Dr. Guacci |
| YGS211 | *MATa smc1-259 ade2-1 trp1-1 can1-100 leu2-3,112 his3-11,15 ura3 GAL psi+ (2μm-TRP)* | This study |
| YGS212 | *MATa smc1-259 ade2-1 trp1-1 can1-100 leu2-3,112 his3-11,15 ura3 GAL psi+ (2μm TRP1 MCD1) (Isolate 1)* | This study |
| YGS213 | *MATa smc1-259 ade2-1 trp1-1 can1-100 leu2-3,112 his3-11,15 ura3 GAL psi+ (2μm TRP1 MCD1) (Isolate 2)* | This study |
| YGS329 | *MATa ura3-52 lys2-801 ade2-101 trp1-Δ63 his3-Δ200 leu2-Δ1 ctf7Δ1::HIS3 ctf7-203:LEU2 + (2μm-TRP1)* | This study |
| YGS330 | *MATa ura3-52 lys2-801 ade2-101 trp1-Δ63 his3-Δ200 leu2-Δ1 ctf7Δ1::HIS3 ctf7-203:LEU2 + (2μm TRP1 MCD1) (Isolate A)* | This study |
| YGS331 | *MATa ura3-52 lys2-801 ade2-101 trp1-Δ63 his3-Δ200 leu2-Δ1 ctf7Δ1::HIS3 ctf7-203:LEU2 + (2μm TRP1 MCD1) (Isolate B)* | This study |
| YGS335 | *MATa NET1-GFP:kanMX6 ade2-1 trp1-1 can1-100 leu2-3,112 his3-11,15 ura3 GAL psi+ (2μm-TRP1)* | This study |

| | | |
|---|---|---|
| YGS337 | *MATa NET1-GFP:kanMX6 ade2-1 trp1-1 can1-100 leu2-3,112 his3-11,15 ura3 GAL psi+ (2µm TRP1 MCD1)* | This study |
| YGS338 | *MATa smc1-259 NET1-GFP:kanMX6 ade2-1 trp1-1 can1-100 leu2-3,112 his3-11,15 ura3 GAL psi+ (2µm-TRP1)* | This study |
| YGS341 | *MATa smc1-259 NET1-GFP:kanMX6 ade2-1 trp1-1 can1-100 leu2-3,112 his3-11,15 ura3 GAL psi+ (2µm TRP1 MCD1)* | This study |
| YBS1042 | *MATα ade2 trp1 his3 leu2::LEU2tetR-GFP ura3::3xURA3tetO112 PDS1-13MYC:TRP1* | Kenna and Skibbens, 2003 |
| YGS333 | *MATa ade2 trp1 leu2::LEU2tetR-GFP ura3::3xURAtetO112 trp1 + (2µm-TRP1) (spore A4B)* | This study |
| YGS334 | *MATa ade2 trp1 leu2::LEU2tetR-GFP ura3::3xURAtetO112 trp1 + (2µm TRP1 MCD1) (spore A4B)* | This study |
| YGS321 | *MATa smc1-259 ade2 trp1 leu2::LEU2tetR-GFP ura3::3xURAtetO112 trp1 + (2µm-TRP1) (sporeE1B)* | This study |
| YGS323 | *MATa smc1-259 ade2 trp1 leu2::LEU2tetR-GFP ura3::3xURAtetO112 trp1 + (2µm TRP1 MCD1) (sporeE1B)* | This study |
| 3327-9B | *Mata scc3-6 GFPLacI-HIS3 his3-11,15 trp1-1 leu2-3,112 ura3-52 bar1 GAL+* | Gift from Dr. Guacci |
| YBS4558 | *MATa ura3-52 lys2-801_amber ade2-101_ochre trp1-Δ63 his3-Δ200 leu2-Δ1 + (2µm URA3)* | This study |
| YBS4562 | *MATa ura3-52 lys2-801_amber ade2-101_ochre trp1-Δ63 his3-Δ200 leu2-Δ1 + (2µm URA3 MCD1)* | This study |
| YBS4567 | *Mata scc3-6 GFPLacI-HIS3 his3-11,15 trp1-1 leu2-3,112 ura3-52 bar1 GAL+ + (2µm URA3)* | This study |
| YBS4568 | *Mata scc3-6 GFPLacI-HIS3 his3-11,15 trp1-1 leu2-3,112 ura3-52 bar1 GAL+ + (2µm URA3 MCD1)* | This study |
| YBS4569 | *Mata scc3-6 GFPLacI-HIS3 his3-11,15 trp1-1 leu2-3,112 ura3-52 bar1 GAL+ + (2µm URA3 MCD1)* | This study |
| KT048 | *MATa scc2-4 MCD1-3HA:TRP1 ade2-1 his3-11,15 leu2-3,112 trp1-1 ura3-1 can1-100* | Tong and Skibbens, 2014 |
| KT046 | *MATa MCD1-3HA:TRP1 ade2-1 his3-11,15 leu2-3,112 trp1-1 ura3-1 can1-100* | Tong and Skibbens, 2014 |
| YGS216 | *MATa MCD1-3HA:TRP1 ade2-1 his3-11,15 leu2-3,112 trp1-1 ura3-1 can1-100 + (2µm LEU2)* | This study |
| YGS217 | *MATa MCD1-3HA:TRP1 ade2-1 his3-11,15 leu2-3,112 trp1-1 ura3-1 can1-100 + (2µm LEU2 MCD1)* | This study |
| YGS218 | *MATa scc2-4 MCD1-3HA:TRP1 ade2-1 his3-11,15 leu2-3,112 trp1-1 ura3-1 can1-100 + (2µm LEU2)* | This study |

| | | |
|---|---|---|
| YGS219 | *MATa scc2-4 MCD1-3HA:TRP1 ade2-1 his3-11,15 leu2-3,112 trp1-1 ura3-1 can1-100 + (2µm LEU2 MCD1) (Isolate 1)* | This study |
| YGS220 | *MATa scc2-4 MCD1-3HA:TRP1 ade2-1 his3-11,15 leu2-3,112 trp1-1 ura3-1 can1-100 + (2µm LEU2 MCD1) (Isolate 2)* | This study |
| BB2788 | *MATa esp1-1 ura3 leu2 can1-100 lys2* | Gift from Dr. Guacci |
| YGS344 | *MATa esp1-1:kanMX6 ura3-52 lys2-801_amber ade2-101_ochre trp1-Δ63 his3-Δ200 leu2-Δ1* | This study |
| YGS348 | *MATa esp1-1:kanMX6 ura3-52 lys2-801_amber ade2-101_ochre trp1-Δ63 his3-Δ200 leu2-Δ1* | This study |
| **YBS255** | *MATa CTF7:LEU2 ctf7::HIS3 ade2-101 his3Δ200 leu2Δ1 lys2-801 trp1Δ63 ura3-53* | Skibbens, 1999 |
| **YMM828** | *MATalpha cft7::HIS3 rad61::URA3 ade2-101 his3Δ200 leu2Δ1 lys2-801 trp1Δ63 ura3-52* | Maradeo and Skibbens, 2010 |
| **YBS4067** | *MATa CTF7:LEU2 ctf7::HIS3 ade2-101 his3Δ200 leu2Δ1 lys2-801 trp1Δ63 ura3-53 + (2µm TRP1)* | This study |
| **YBS4071** | *MATa CTF7:LEU2 ctf7::HIS3 ade2-101 his3Δ200 leu2Δ1 lys2-801 trp1Δ63 ura3-53 + pGS2 (2µm TRP1 PDS5)* | This study |
| **YBS4069** | *MATalpha cft7::HIS3 rad61::URA3 ade2-101 his3Δ200 leu2Δ1 lys2-801 trp1Δ63 ura3-52 + (2µm TRP1)* | This study |
| **YBS4075** | *MATalpha cft7::HIS3 rad61::URA3 ade2-101 his3Δ200 leu2Δ1 lys2-801 trp1Δ63 ura3-52 + pGS2 (2µm TRP1 PDS5) iso A* | This study |
| **YBS4076** | *MATalpha cft7::HIS3 rad61::URA3 ade2-101 his3Δ200 leu2Δ1 lys2-801 trp1Δ63 ura3-52 + pGS2 (2µm TRP PDS5) iso B* | This study |
| **YBS2031/ KT046** | *MATa ade2-1 his3-11,15 leu2-3,112 trp1-1 ura3-1 can1-100 MCD1-3HA:TRP1* | Tong and Skibbens, 2014 |
| **YBS2033/ KT048** | *MATa ade2-1 his3-11,15 leu2-3,112 trp1-1 ura3-1 can1-100 scc2-4 MCD1-3HA:TRP1* | Tong and Skibbens, 2014 |
| **YBS4864** | *MATα eco1::HIS3 rad61::URA3 esp1-1:kanMX6 ade2-101 his3Δ200 leu2Δ1 lys2-801 trp1Δ63 ura3-52 (spore 10A)* | This study |
| **YBS4862** | *MATa esp1-1:kanMX6 ura3-52 lys2-801_amber ade2-101_ochre trp1-Δ63 his3-Δ200 leu2-Δ1 (spore 7C)* | This study |
| **YBS4863** | *MAT esp1-1:kanMX6 ura3-52 lys2-801_amber ade2-101_ochre trp1-Δ63 his3-Δ200 leu2-Δ1 (spore 3B)* | This study |
| **YBS4860** | *MATa eco1::HIS3 rad61::URA3 ade2-101 his3Δ200 leu2Δ1 lys2-801 trp1Δ63 ura3-52 (spore 4C)* | This study |
| **YBS4861** | *MATα eco1::HIS3 rad61::URA3 ade2-101 his3Δ200 leu2Δ1 lys2-801 trp1Δ63 ura3-52 (spore 9D)* | This study |

| | | |
|---|---|---|
| **YGS375** | *MATalpha eco1::HIS3 rad61::URA3 pdr5::TRP ade2-101 his3Δ200 leu2Δ1 lys2-801 trp1Δ63 ura3-52* | This study |
| **YGS377** | *MATa eco1::HIS3 rad61::URA3 pds5::TRP ade2-101 his3Δ200 leu2Δ1 lys2-801 trp1Δ63 ura3-52* | This study |
| **YGS352** | *MATa ubr1::kanMX6 ura3-52 lys2-801_amber ade2-101_ochre trp1-Δ63 his3-Δ200 leu2-Δ1 (Isolate c1)* | This study |
| **YGS354** | *MATalpha ubr1::kanMX6 eco1::HIS3 rad61::URA3 ade2-101 his3Δ200 leu2Δ1 lys2-801 trp1Δ63 ura3-52 (Isolate e1)* | This study |
| **YGS355** | *MATalpha ubr1::kanMX6 eco1::HIS3 rad61::URA3 ade2-101 his3Δ200 leu2Δ1 lys2-801 trp1Δ63 ura3-52 (Isolate e2)* | This study |
