## Supplementary material for "Aberrant cohesin function in *Saccharomyces cerevisiae* activates Mcd1 degradation to promote cell lethality": S2 Table

| Name (oRVS) | DNA oligos Sequence 5' to 3' |
|---|---|
| **8** | ACG ACT CAT CTC CAT GCA GTT |
| **23** | GAT TGT CGC ACC TGA TTG CC |
| **3370** | AGG ACT TTG CAA CTG AAG CAG CAG ATT TGA GAT ATA TTC TGG GGA ACA AAA GAA GTA TTA CGG ATC CCC GGG TTA ATT AA |
| **3371** | ATA GTC CTG CTT TTA TCA ATT ATT TGT AAA ACT GCG AGA TTA CTG TTA GTG TTG TAT GGT GAATTC GAG CTC GTT TAA AC |
| **3372** | GTC CAA CAG ATA TGC GTA AAG |
| **3373** | CAG AAA TTG CTG ATT TTT ACT CCT ACT TAG TAT ACA TTT CAC TAA ACA ATA CGT TTT ACC CGGATC CCC GGG TTA ATT AA |
| **3374** | AGT TTT CAG GTA TCT AAG ATA AAA ATA TAT GGT AAA TAC CTT TAA CGA ATA TTA TAA AAT GAATTC GAG CTC GTT TAA AC |
| **3375** | TTG CTT ACT TGT ATC GCT ATT |
| **3376** | CCG TGC AAA ATA TCC AGG ACG TCT ATA CAC AGT GTT TAC AAC TCA GCT TAT ATT CAT ATC CGG ATC CCC GGGTTA ATT AA |
| **3377** | CTA GCT TAA AAA ATG CGT TGA ATA TAT ATT ATT AAA TAT ATA TAT TTG AAG GGG AGT TGA GAA TTC GAG CTCGTT TAA AC |
| **3378** | TGT TCA CCA GAG GAT TCT TTC |
| **3379** | TTC TCC CTT TTT TCC CCT TTG TTT TCT CTC ATA GTC TTG TAA CCT CAG CTT TTG TTC ATT CGG ATC CCC GGG TTA ATT AA |
| **3380** | AAA TGG ATG ACT GCC AAT AGG ACA TAT TTT CAT ATT AAC ATA CTT CAG AAG CGG TAT TGT GAA TTC GAG CTC GTT TAA AC |
| **3381** | ATA CTT CGA CGC AAA AAG CCG |
| **3385** | AGA GAT AGA AAG GGC TTT CAC CGT TTT TAT GCT AAT CGT GCT AGC TGA TAA TAA TCA GAT CGG ATC CCC GGG TTA ATT AA |
| **3386** | TAT GTA TAT GTA TGT GGA GGA TAT AAC ACA AAC AGT GGA AAA GTG GTA GAA TAA TTA GTA GAA TTC GAG CTC GTT TAA AC |
| **3387** | CTG GTG AAA TTC TGA GAT CGT |
| **1203** | TTT AGG TAA GAA GAA GAA GCC AAG TGG TGG ATT TGC ATC ATT AAT AAA AGA TTT CAA GAA AAA ACG GAT CCC CGG GTT AAT TAA |
| **1204** | TGC TTG ATT ATT TTT TTT TAC TAG CTT TCT GTG ACG TGT ATT CTA CTG AGA CTT TCT GGT ATC AGA ATT CGA GCT CGT TTA AAC |
| **3390** | CTT GTT TCA AGA GGC ATC CCA |
| **3346** | GGGAAGAAGGAACAAGACAAA (qPCR *MCD1* sense) |
| **3347** | TCCCTAGATCCCAACCAATAG (qPCR *MCD1* antisense) |
| **3348** | CAGGCAATGTCACGGATAG (qPCR *ALG9* sense) |
| **3349** | CCTTCACACCACCTTGATTTA (qPCR *ALG9* antisense) |
